# Drugs targeting microRNAs that regulate DNA damage sensing and repair mechanisms: *A computational drug discovery approach*

**DOI:** 10.1101/2024.06.18.599590

**Authors:** S.B Ravindranath, S. Vratin, B.S. Lakshmi, D.C. Ramirez, S.E. Gomez Mejiba

**Affiliations:** Department of Biotechnology, Manipal Institute of Technology, Manipal, Manipal Academy of Higher Education, Manipal 576104, India; Department of Biological Sciences, Carnegie Mellon University, 4400 5th Ave, Rm 255 Mellon Institute, Pittsburgh, PA15213, USA; Department of Science and Humanities, PES university, BSK III stage, Bengaluru 560085. India; Laboratory of Experimental and Translational Medicine, National Bureau of Science and Technology-San Luis, National University of San Luis, San Luis, San Luis 5700, Argentina; Laboratory of Nutrition and Experimental Therapeutics, National Bureau of Science and Technology-San Luis, National University of San Luis, San Luis, San Luis 5700, Argentina

**Author notes:** **Corresponding Authors:** Ravindranath, S.B.; Ramirez, DC; and Gomez Mejiba, SE. Authors’ e-mail and ORCID: Ravindranath, S Bilachi ORCID: 0000-0002-3713-3646; Vratin Srivastava ORCID: 0000-0001-9341-2570; Lakshmi B. Sanganabasappa ORCID: 0009-0005-4300-1534; Ramirez, Dario C. ORCID: 0000-0001-6725-3326; Gomez Mejiba, Sandra E. ORCID: 0000-0002-8515-0483.

**Keywords:** DNA damage sensing, DNA damage repair, gene expression, miR, computational drug discovery, gene-disease-miR-drug network

## Abstract

DNA damage-sensing (DDS) and DNA damage-repair (DDR) mechanisms are essential for the fidelity of genetic information transmission. Failure to accomplish an effective DDS/DDR mechanism can lead to cell death or otherwise to cell transformation and cancer development. microRNAs (miRs) are short noncoding RNAs that primarily function as micromanagers of gene expression. Herein, we aimed to investigate the links between miRs and the translation of specific mRNAs encoding proteins involved in genomic DDS and DDR and to screen drugs that have high binding affinity to the selected miRs, which may serve as cancer therapeutics. To accomplish these aims, we used a variety of computational methods spanning data analysis, molecular modeling, and simulation tools (*i.e.,* PyRx, Biopython, ViennaRNA, RNAComposer, AutoDock Vina, OpenBabel, PyMOL, Discovery Studio, MarvinSketch). The genes and miRNAs involved in the DDS and DDR mechanisms were retrieved from either the literature or various online databases (*e.g.,* miRDB). miR data were further cleaned and prepared using scripts, and various libraries were used to obtain their 3-D structures. Genes interacting with miRs were enriched based on multiple database annotations using Enrichr KG. Then, we used docking analyses to virtually screen compounds to serve as ligands for the miRs. Finally, we generated gene-disease-miR-drug networks to study the linkages between the compounds and miR molecules under investigation. For the first time, we were able to identify five compounds that could be repurposed for downregulating miRs that are linked to inhibition of translation of mRNA involved in the DDS and DDR processes. The gathered candidate drugs can be useful for preventing cell transformation and cancer development.

## 1. INTRODUCTION

Spontaneous and induced DNA polymerase errors or environmentally induced genomic damage must be sensed and repaired by well-regulated DNA direct and indirect repair mechanisms. The DNA damage response is sensed by specific proteins, leading to the activation of signaling pathways to engage repair processes (Harper & Elledge, 2007). If genomic DNA damage is not repaired, the cell can undergo apoptosis or, in the worst case, cell transformation and tumor development (Chatterjee & Walker, 2017). Several studies have shown that in most cancers, alterations to the DNA damage-sensing (DDS) and -response (DDR) pathways occur (Bouwman & Jonkers, 2012; Ghosal & Chen, 2013).

When there is no significant distortion to the DNA helix due to DNA damage, the base excision repair (BER) process is activated. This process has been shown to occur in the G1 phase of the cell cycle (Dianov & Hübscher, 2013). However, when the damage to the DNA is bulky and is responsible for twisting the DNA helix, mostly due to damage from UV radiation or chemotherapeutic agents, the nucleotide excision repair (NER) process is activated (Masutani et al., 1994; Nishi et al., 2005). When simple incorrect Watson-Crick base pairing (mismatches) or strand slippage occurs, the mismatch repair (MMR) mechanism is activated (Bindra & Glazer, 2007; Mihaylova et al., 2003; Nakamura et al., 2008). Multiple other repair mechanisms also exist, such as interstrand cross-link repair, translesion synthesis, and single- and double-strand break repairs (Chatterjee & Walker, 2017). Therefore, it is essential to identify genes encoding DDS and DDR proteins to ensure efficient repair mechanisms.

Small, noncoding RNAs called microRNAs (or miRs) play a key role in posttranscriptional gene regulation by silencing gene expression at the level of translation. These micromanagers of gene expression are approximately 18–23 nucleotides in length (Felekkis et al., 2010). In mammals, miRs regulate approximately 30% of all protein-encoding genes (Filipowicz et al., 2008; Kozomara & Griffiths-Jones, 2011). miRs are transcribed as long transcripts by RNA polymerase II from their encoding genes as long transcripts named pri-miRs. Then, pre-miRNAs are produced inside the nucleus by the cleavage of the pri-miRNA by the Drosha RNase III endonuclease and the DCGR8 enzyme (Lee et al., 2002). The miR-miR* complex is an incomplete double-stranded molecule, where miR is the mature miR and miR* is the opposing arm (Aravin et al., 2003; Lagos-Quintana et al., 2002; Vishnoi & Rani, 2017). The miR-miR* complex is loaded into the RNA-induced silencing complex (RISC), which contains the argonaut (AGO) protein–the catalytic subunit (Lodish et al., 2008). This RISC complex then binds to the target mRNA and leads to translation suppression (Sanghvi & Steel, 2011; Vishnoi & Rani, 2017).

Herein, we aimed to investigate the links between miRs and the translation of specific mRNAs encoding proteins involved in genomic DDS and DDR processes. Using this information, we then screened those drugs that had the highest binding affinity to the selected miRs, which may serve as novel cancer therapeutics.

## 2. METHODS

### 2.2. Computational workflow

Various computational methods have been used to extract, clean, and standardize the data we wish to analyze and simulate. **Fig 1** shows the workflow used to find DDS/DDR-modulating miR sequences, obtain the 3-D structure of those miRs, and identify potential ligands to inhibit their action. Initially, the list of genes that are involved in DDR processes was collected through an extensive literature search and collated into one list of target genes (Aravind et al., 1999; Eisen & Hanawalt, 1999; Knijnenburg et al., 2018; Lange et al., 2011; Montelone, 2006; Ronen & Glickman, 2001; Wood et al., 2001).

**Figure 1.**
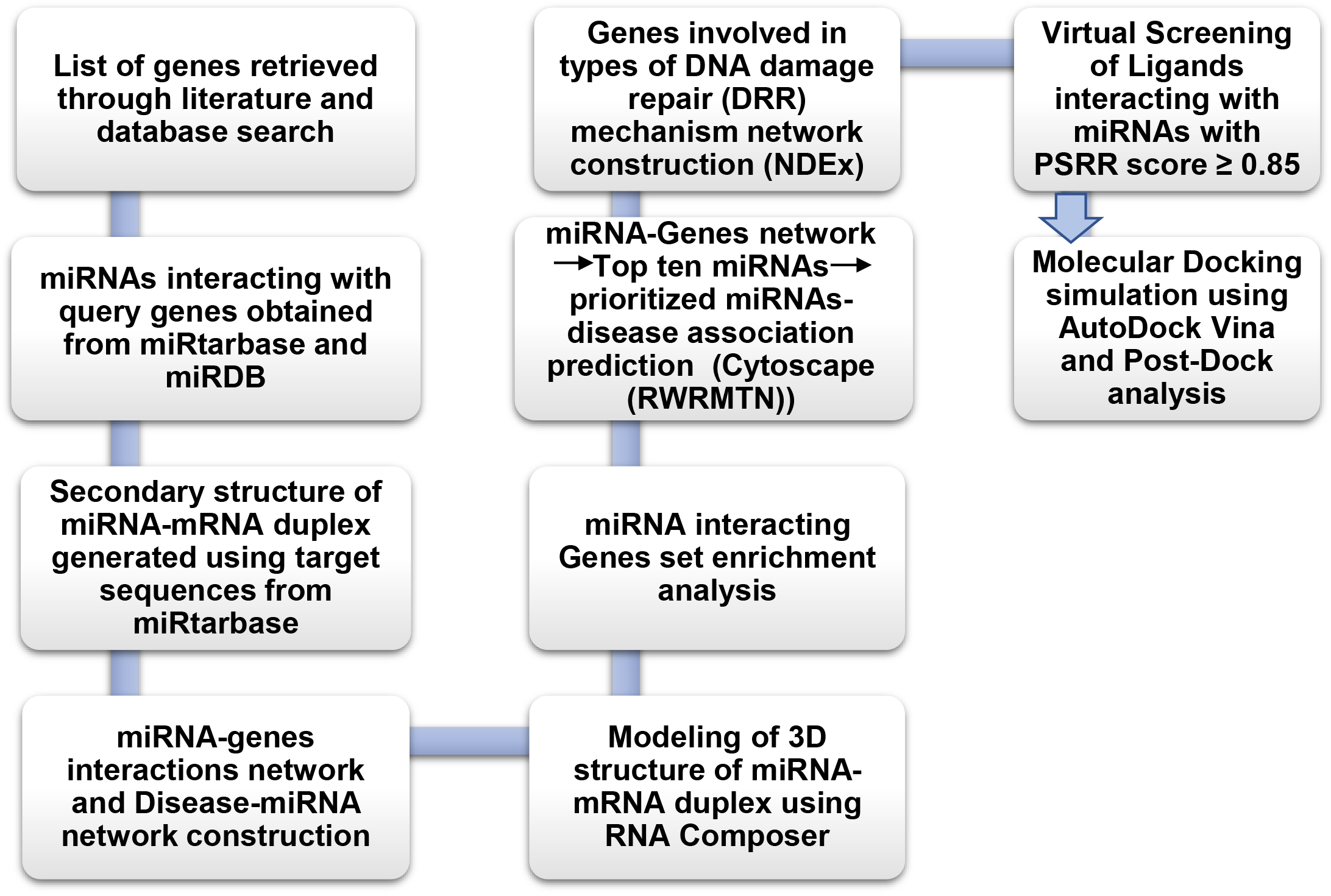
Workflow of the methodology

### 2.2. miR Data

Multiple database sources contain data regarding both experimentally verified and predicted targets of miRs. We obtained a list of miRs from both sources. For experimentally verified targets, we used miRTarBase (H. Y. Huang et al., 2020), which only contains target data that have been experimentally validated. From the data obtained, we filtered out the desired miRs by cross-referencing the gene targets with the initial gene list we had procured. We obtained 32 unique miRs through this procedure, some of which target multiple genes from our list. To retrieve miRs that have our desired genes as predicted targets, we used the database miRDB (Y. Chen & Wang, 2020). This database uses a proprietary prediction algorithm named miRTarget (Liu & Wang, 2019) to detect possible mRNA targets for miRs. We were able to obtain 83 miRs targeting DDR and DDS-related transcripts through this database.

Then, we obtained the 3-D structures of miR-mRNA duplexes by searching databases, or in the absence of X-ray data, there exist multiple algorithms and libraries for predicting the structure of these complexes that may aid in our research. To obtain the 3-D structure, it was necessary to first find the sequences and secondary structures.

### 2.3. miR sequences

To obtain the miRNA sequences, we first downloaded a database of all known miRs acting on human genes from miRBase (Griffiths-Jones, 2004; Griffiths-Jones et al., 2006, 2008; Kozomara et al., 2019; Kozomara & Griffiths-Jones, 2011, 2014). miRBase is a virtual database of published miR sequences and annotations. Then, to retrieve the sequences for the selected miRs, we used a Python script that was written using the Biopython library (Chapman & Chang, 2000; Cock et al., 2009). The Biopython library allowed efficient parsing of FASTA files and made it easy to write into another FASTA file.

### 2.4. miR-mRNA duplex secondary and 3D tertiary structure

RNAs are very similar to proteins in the sense that they also have secondary and tertiary structures based on the way they fold owing to noncovalent bonding and interaction between different functional groups within their nucleotide chains. Before predicting the 3D tertiary structure, we deciphered the secondary structure from the miR-mRNA sequence. The gene binding location (miR-target RNA sequence) was obtained from miRTarBase.

A molecule’s most thermodynamically stable structure is the one with the lowest molecular free energy (MFE) (Ronny et al., 2016). The MFE structure was thus one of the first goals of our structure prediction strategy (Irmtraud & Meyer, 2007). Any sequence has a finite number of valid secondary structures. In theory, the MFE structure can be derived by computing the free energy for each potential base-pairing pattern using the experimentally discovered set of energy rules (Higgs, 2000). Herein, we used ViennaRNA to determine secondary structures (Lorenz et al., 2011). For the secondary structure of miRs, the notation used is as suggested by Zhong et al., where a base is represented by one character. A base is paired with another base ahead of it if the parenthesis is open. A base with closed parentheses is paired with another base behind it. Periods, or dots, denote an unpaired base. Therefore, there will always be an equal number of open and closed parentheses (Zhong et al., 2013).

To obtain the predicted tertiary 3D structures of the miR-mRNA, the tool RNAComposer was used. This tool’s workflow entails breaking down the user-defined RNA secondary structure into its parts. According to Popenda et al. (2010), this fragmentation algorithm provides stems, loops (apical, bulge, internal, and n-way junctions), and single strands that are all closed by canonical base pairs (s). This serves as the input for automatically searching possible related tertiary structure elements in the RNA FRABASE dictionary (Popenda et al., 2010, 2012). Finally, we used energy reduction in torsion angle space and Cartesian coordinates to refine the generated structure to create the final, high-quality RNA 3D model.

### 2.5. Construction of networks for the interaction miR target genes

The roles of miRs in diverse regulatory signaling pathways are well established. In this respect, it is important to identify target genes specific to miRs to gain insights into the regulatory molecular mechanism and their role in causing diseases. Target identification through *in vitro* and *in vivo* methods is resource intensive and expensive. Therefore, the computational prediction of the mRNAs interacting with miRs is a significant approach to understanding miR-target gene interactions. These computational data need to be further validated considering the prioritized miR-mRNA interactions by using experimental methods. Network Data Exchange Integrated Query (NDEx IQuery) leverages building networks based on gene sets from pathways. It specifically integrates resources from curated pathways (WikiPathways & SIGNOR) and integrates them with Cytoscape, leading to archival and sharing of the analyzed results. The overlapping genes were predicted to be involved in multiple DDR mechanisms to build networks based on the INDRA GO system.

### 2.6. miR-disease network construction

Because miRs are involved in regulating gene expression, they are highly associated with the pathophysiology of multiple diseases, including cancers. The experimental approach to derive the relations of miRs to disease is also laborious and expensive. *In silico* prediction of miR-disease interactions can lead to the discovery of miR functions as biomarkers and therapeutics. miRNet has effectively built miR-disease networks by integrating database information. This platform is composed of the input data and currently available knowledge base on miR functions. Prediction of the betweenness of the miRs for the prioritization of the miR-disease correlations was adopted (Chang et al., 2020). It provides high-performance miR-centric networks for visual data analysis by integrating fourteen varied miR databases for six host organisms, including humans.

### 2.7. miR interacting gene set enrichment prediction

RNA expression analysis is a crucial tool in bioinformatics for obtaining novel insights into biological processes, diseases, signaling pathways, drug targets, and cell types. This information is significant for leading to knowledge-based discovery of gene potential. Gene set enrichment analysis is a powerful analytical technique that provides novel insights into gene involvement in the abovementioned areas. We used Enrichr KG as a unique tool for the integration of the key insights of biological information about mRNA and cancer. Gene enrichment is a crucial step in multiomics data analysis based on experimental results. Enrichr has gained popularity as a gene enrichment analysis web server by integrating gene set enrichment predictions across various knowledge bases (domains and libraries).

Enrichr KG is a web application with knowledge graph applications for the integration of gene set enrichment analysis and interactive visual outputs. It provides subgraphs including nodes and edges by connecting the genes with the enriched features. Therefore, we generated an info-graphical output by combining the results across 26 libraries with the enriched gene set. It orients the significant associations masked between the genes and annotated enriched data from multiple knowledge resources based on multiple categories, such as ontologies, pathways, cells, transcription factors, and diseases/drugs.

### 2.8 Virtual Screening of Ligands

To identify ligands that not only interact with the miR molecules of interest but also downregulate them, we used a web server named PSRR (Yu et al., 2022). This server uses the random forest algorithm to predict up- and downregulation pairs for miR-small molecules. The downregulated ligands were obtained by entering the respective sequences to be targeted. Ligands with a threshold score >0.85 were selected, where PSRR predictions were based on random forest algorithms. Furthermore, the machine learning model was trained on empirical data. (https://rnadrug.shinyapps.io/PSRR/). When the PSRR server downregulation ligand-miR model’s prediction score was >0.41 (suggestion rate), the small molecules were significant for further study. These ligands were further screened for ADMET properties using SWISSADME and considering multiple factors. Finally, the respective ligands were converted to the desired file extension using Open Babel (O’Boyle et al., 2011).

### 2.9 Molecular docking and analysis

Docking between selected small molecule ligands and miR-mRNA duplexes was performed using AutoDock Vina (Trott & Olson, 2009). For further analysis, we selected the conformations with the highest (most negative) binding energy. Interactions were visualized on PyMOL (https://pymol.org/2/). Nucleotides participating in hydrogen bonding and other interactions with the duplexes were studied and analyzed using BIOVIA Discovery Studio 3.5 (https://www.3ds.com/products-services/biovia/products/molecular-modeling-simulation/biovia-discovery-studio/).

## 3. RESULTS

### 3.1. Gene List

We were able to obtain data for 48 genes along with their specific functions using the DAVID functional annotation tool, as shown in **Table 1**. The workflow for the prediction of the highest ranked miRs, mRNAs, and the association of miR-gene, disease-miRs, genes-DDR mechanism, and the docking results is shown in **Fig 1**.

**Table 1:** List of genes involved in DDS/DDR processes and their specific functions

| Sl. No. | Gene | Selected Functions | PubMed Reference ID |
| --- | --- | --- | --- |
| 1. | ERCC8 | cellular response to DNA damage stimulus | 11782547 |
| 2. | POLD1 | DNA synthesis involved in DNA repair | 1730053 |
| 3. | ERCC1 | double-strand break repair via nonhomologous end joining | 14690602 |
| 4. | CCNH | transcription initiation from RNA polymerase II promoter | 7533895 |
| 5. | XPA | nucleotide-excision repair, DNA incision | 21148310 |
| 6. | CHD1L | DNA repair | 19661379 |
| 7. | ERCC4 | nucleotide-excision repair | 8797827 |
| 8. | DDB2 | UV-damage excision repair | 14751237 |
| 9. | SUMO1 | positive regulation of proteasomal ubiquitin-dependent protein catabolic process | 18408734 |
| 10. | XPC | UV-damage excision repair | 8077226 |
| 11. | ACTB | regulation of cell cycle | 27153538 |
| 12. | PCNA | mismatch repair | 11005803 |
| 13. | CUL4B | ribosome biogenesis | 26711351 |
| 14. | ERCC3 | positive regulation of apoptotic process | 16914395 |
| 15. | ERCC6 | transcription by RNA polymerase II | 7664335 |
| 16. | PARP1 | apoptotic process | 15565177 |
| 17. | ERCC5 | nucleotide-excision repair | 11259578 |
| 18. | EP300 | positive regulation of transcription, DNA-templated | 27256286 |
| 19. | ERCC2 | chromosome segregation | 20797633 |
| 20. | WRN | cellular response to DNA damage stimulus | 18203716 |
| 21. | POLE | DNA-dependent DNA replication | 33051204 |
| 22. | BLM | DNA repair | 7585968 |
| 23. | RAD51C | DNA repair | 19451272 |
| 24. | ATR | cellular response to DNA damage stimulus | 15538388 |
| 25. | RBBP8 | DNA double-strand break processing involved in repair via single-strand annealing | 18716619 |
| 26. | ATM | protein phosphorylation | 17694070 |
| 27. | BRCA1 | response to ionizing radiation | 17525340 |
| **28.** | **BRCA2** | double-strand break repair via homologous recombination | 21719596 |
| **29.** | **NBN** | mitotic G2 DNA damage checkpoint signaling | 11438675 |
| **30.** | **RAD51D** | telomere maintenance | 15109494 |
| **31.** | **MRE11** | cellular response to DNA damage stimulus | 29670289 |
| **32.** | **RAD50** | telomere maintenance | 10888888 |
| **33.** | **POLH** | regulation of DNA repair | 10398605 |
| **34.** | **BRIP1** | DNA damage checkpoint signaling | 14576433 |
| **35.** | **RTEL1** | regulation of double-strand break repair via homologous recombination | 18957201 |
| **36.** | **BARD1** | cellular response to DNA damage stimulus | 15905410 |
| **37.** | **ABL1** | DNA damage induced protein phosphorylation | 18280240 |
| **38.** | **UBE2T** | DNA repair | 16916645 |
| **39.** | **FANCF** | protein monoubiquitination | 24910428 |
| **40.** | **FANCA** | protein-containing complex assembly | 9398857 |
| **41.** | **FANCD2** | response to gamma radiation | 12874027 |
| **42.** | **FAN1** | DNA repair | 20603073 |
| **43.** | **FANCI** | positive regulation of protein ubiquitination | 18029348 |
| **44.** | **FANCC** | DNA repair | 1574115 |
| **45.** | **FANCG** | mitochondrion organization | 17060495 |
| **46.** | **MSH6** | mismatch repair | 8782829 |
| **47.** | **MLH1** | mismatch repair | 23603115 |
| **48.** | **MSH2** | postreplication repair | 7923193 |

### 3.2. Screening for miRs with DDR genes as targets

Among the 84 miRs, we obtained 32 unique miRs that have DDS/DDR genes as targets, and the experimental validation is shown in **Table 2**. Therein, miRs with query genes as predicted targets are shown along with their miRTarget score. For docking studies, miRs targeting more than 1 unique gene were selected.

**Table 2:**
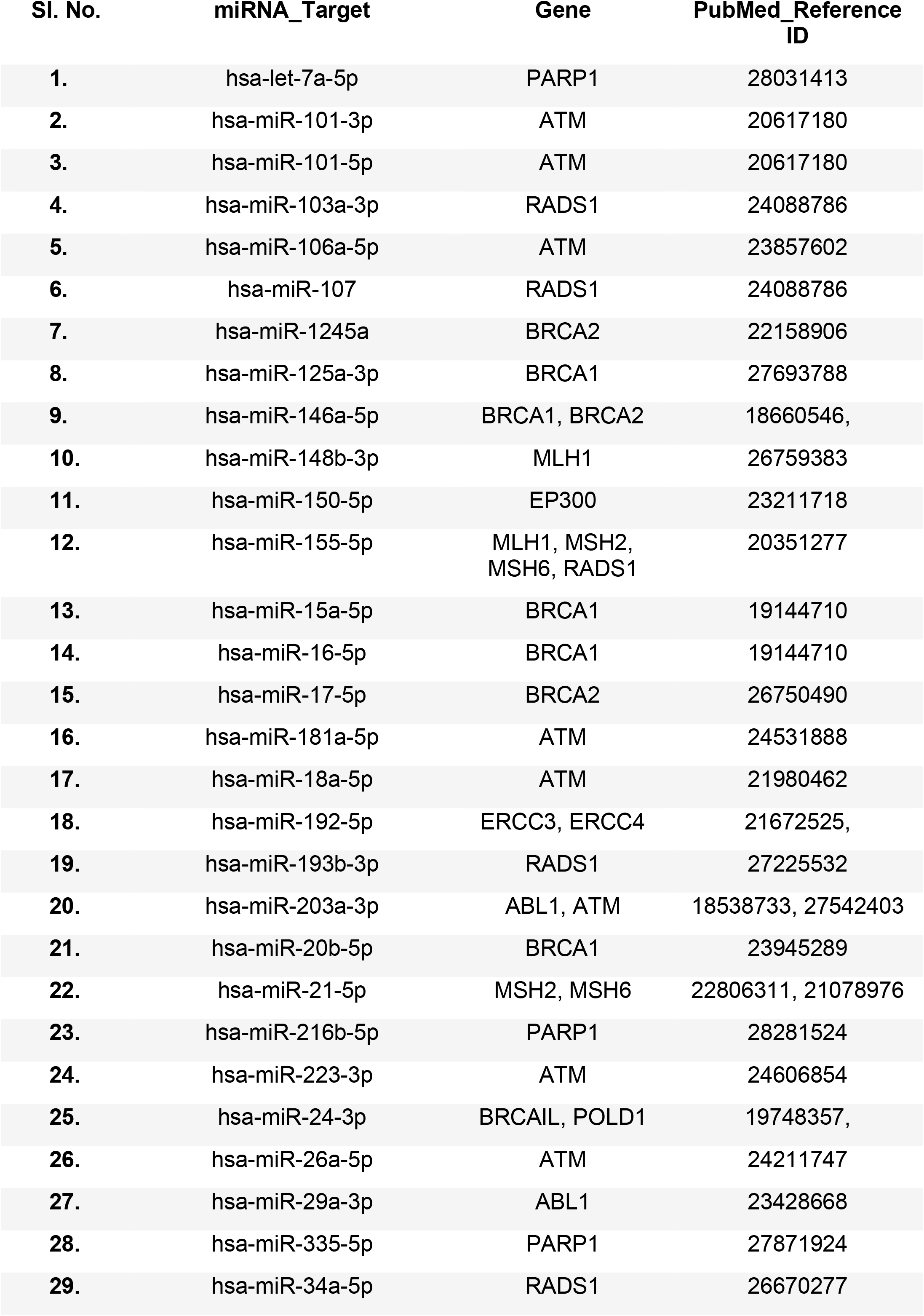
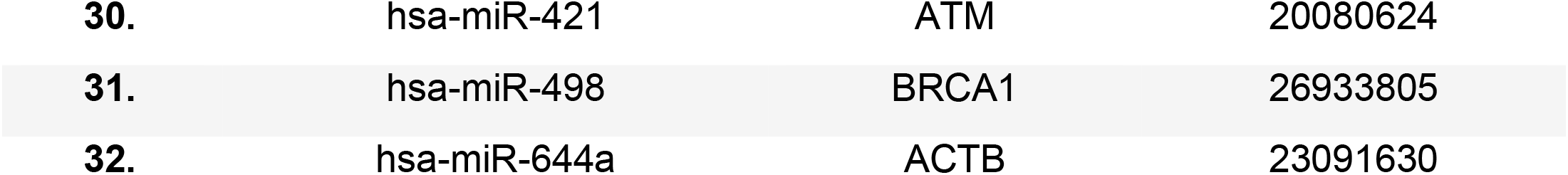
Unique miRs targeting queried DDS/DDR genes set with PubMed IDs

| Sl. No. | miRNA_Target | Gene | PubMed_Reference ID |
| --- | --- | --- | --- |
| 1. | hsa-let-7a-5p | PARP1 | 28031413 |
| 2. | hsa-miR-101-3p | ATM | 20617180 |
| 3. | hsa-miR-101-5p | ATM | 20617180 |
| 4. | hsa-miR-103a-3p | RADS1 | 24088786 |
| 5. | hsa-miR-106a-5p | ATM | 23857602 |
| 6. | hsa-miR-107 | RADS1 | 24088786 |
| 7. | hsa-miR-1245a | BRCA2 | 22158906 |
| 8. | hsa-miR-125a-3p | BRCA1 | 27693788 |
| 9. | hsa-miR-146a-5p | BRCA1, BRCA2 | 18660546, |
| 10. | hsa-miR-148b-3p | MLH1 | 26759383 |
| 11. | hsa-miR-150-5p | EP300 | 23211718 |
| 12. | hsa-miR-155-5p | MLH1, MSH2,<br>MSH6, RADS1 | 20351277 |
| 13. | hsa-miR-15a-5p | BRCA1 | 19144710 |
| 14. | hsa-miR-16-5p | BRCA1 | 19144710 |
| 15. | hsa-miR-17-5p | BRCA2 | 26750490 |
| 16. | hsa-miR-181a-5p | ATM | 24531888 |
| 17. | hsa-miR-18a-5p | ATM | 21980462 |
| 18. | hsa-miR-192-5p | ERCC3, ERCC4 | 21672525, |
| 19. | hsa-miR-193b-3p | RADS1 | 27225532 |
| 20. | hsa-miR-203a-3p | ABL1, ATM | 18538733, 27542403 |
| 21. | hsa-miR-20b-5p | BRCA1 | 23945289 |
| 22. | hsa-miR-21-5p | MSH2, MSH6 | 22806311, 21078976 |
| 23. | hsa-miR-216b-5p | PARP1 | 28281524 |
| 24. | hsa-miR-223-3p | ATM | 24606854 |
| 25. | hsa-miR-24-3p | BRCA1, POLD1 | 19748357, |
| 26. | hsa-miR-26a-5p | ATM | 24211747 |
| 27. | hsa-miR-29a-3p | ABL1 | 23428668 |
| 28. | hsa-miR-335-5p | PARP1 | 27871924 |
| 29. | hsa-miR-34a-5p | RADS1 | 26670277 |
| 30. | hsa-miR-421 | ATM | 20080624 |
| 31. | hsa-miR-498 | BRCA1 | 26933805 |
| 32. | hsa-miR-644a | ACTB | 23091630 |

### 3.3. miRNA‒mRNA duplex structure

miR-mRNA duplex structures were generated for BRCA1-24-3p, POLD1-24-3p, ABL1-203a-3p, ATM-203a-3p, MSH6-21-5p, ERCC3-192-5p, ERCC4-192-5p, BRCA1-146a-5p, BRCA2-146a-5p, MLH1-155-5p, MSH6-155-5p, and RAD51-155-5p, as shown in **Table 3**. **Supplementary** Figure 2 shows the 2D structure obtained for each of the selected miRs using the RNAfold Web Server. The 3D structures for the queried miR-mRNA complexes are shown in **Figure 2**.

**Figure 2.**
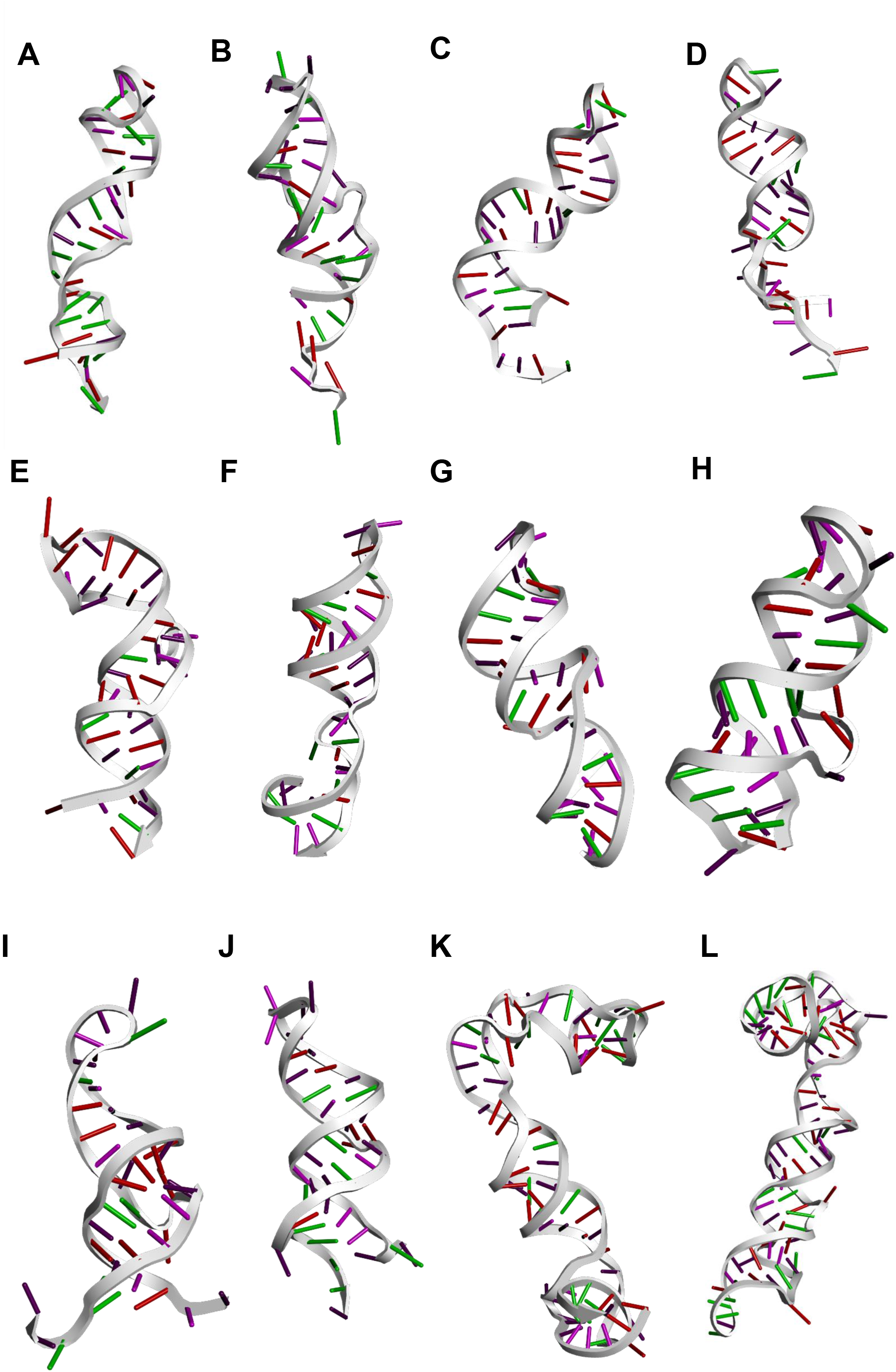
miR-mRNA complexes as modeled 3D structures. RNAComposer data for **A**) BRCA1-24-3p, **B**) POLD1-24-3p, **C**) ABL1-203a-3p, **D**) ATM-203a-3p, **E**) MSH6-21-5p, **F**) ERCC3-192-5p, **G**) ERCC4-192-5p, **H**) BRCA1-146a-5p, **I**) BRCA2- 146a-5p, **J**) MLH1-155-5p, **K**) MSH6-155-5p, and **L**) RAD51-155-5p.

**Table 3:**
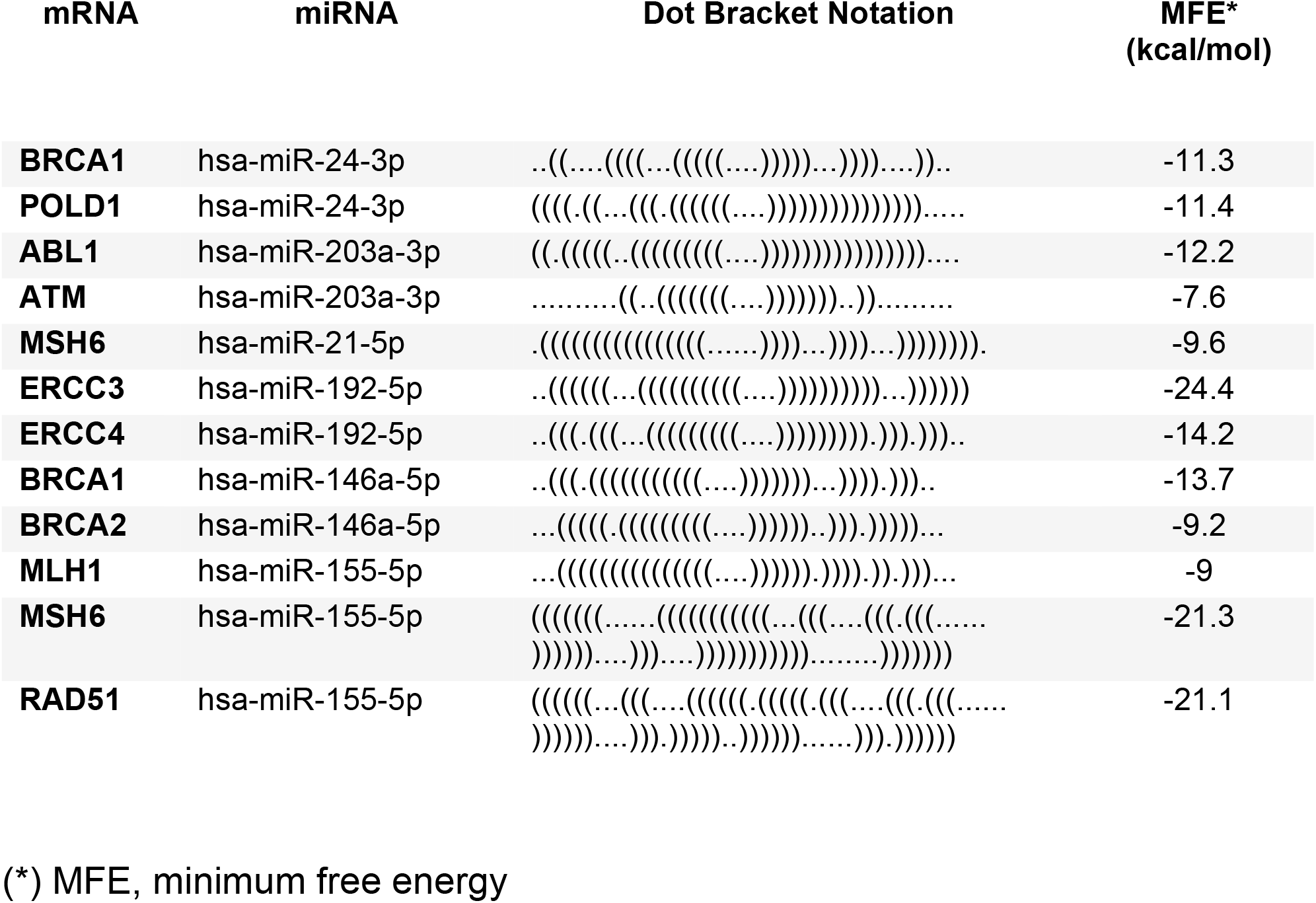
Dot-bracket notation and MFE calculations generated from the RNAfold web server for secondary structures of the mRNA-miR duplexes

### 3.5. Prediction and orientation of query genes involved in DDR mechanisms

Prediction and orientation of the query genes involved in the DDR mechanisms are crucial to prioritize the genes involved in significant steps for developing novel iRNAs as therapeutics. The NDEx IQuery web user ranks and orients the queried genes based on the network overlapping potential and calculates the cumulative distribution based on an adjusted probability score to identify a high false discovery rate. We were able to predict that the queried genes are distributed in 12 networks (**Fig. 3**) involved in specific DDR mechanisms. NDEx IQuery integrates curated pathway information to build gene interaction networks based on annotated Gene Ontology terms (Pillich et al. 2023). This prediction of the genes involved in DDR mechanisms provides novel insights into the functions of specific genes.

**Figure 3.**
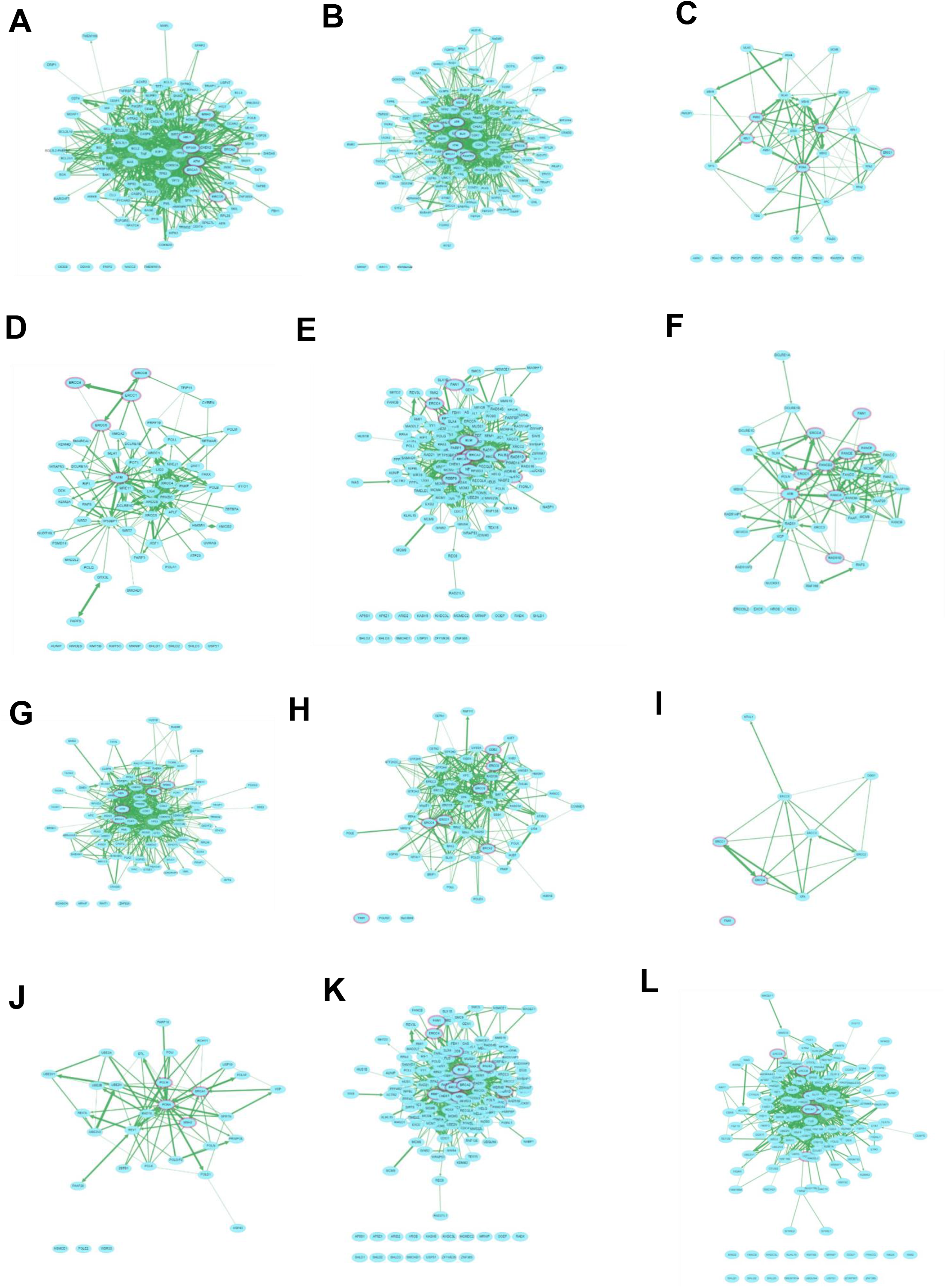
**Oriented genes in networks associated with specific DNA damage repair mechanisms: A**) intrinsic apoptotic signaling pathway response to DNA damage, **B**) DNA damage checkpoint, **C**) mismatch repair, **D**) double-strand break repair (DSBR), **E**) DSBR via homologous recombination, **F**) interstrand cross-link repair, **G**) mitotic DNA integrity checkpoint, **H**) nucleotide excision repair (NER), **I**) NER, DNA incision, **J**) postreplication repair, **K**) telomere maintenance, and **L**) regulation of DNA repair.

### 3.6. Prediction and network analysis of the miR-disease network

As a comprehensive web interface, miRNet (http://www.mirnet.ca/) predicts the target genes for significant disease-miR interactions by integrating the data from eleven miRNA databases (*i.e.,* miRTarBase, TarBase, miRanda, miRecords, miR2Disease, PhenomiR, SM2miR, PharmacomiR, EpimiR, HMDD, and starBase). miRNet 1 was effective in predicting gen-miR interactions. Later, based on transcription factor and single nucleotide polymorphism data affecting the function of miRs, a >5-fold knowledge base enhancement of miR-disease associations was incorporated (see **Suppl. Fig. 1**). Additionally, the visualization of the multifaceted networks for analyzing miRs is enabled. It provided the disease information from the DisGeNET database and ranked them based on the number of hits, P value, and adj. Pval. The diseases associated with high-ranked miRs by miRNet are shown in **Supplementary Table 1**.

Autosomal recessive predisposition was ranked first with 161 hits (**see Suppl. Table 1**). The gathered miR list includes multiple cancer types, multiorganelle-associated diseases, and other disorders. The disease-miR interaction is ordered in ascending order along with the P value and adj. Pval, as shown in **Supplementary Table 1**.

### 3.6. miR interacting gene set enrichment analysis

The graphical representation shown in **Fig. 4A** highlights those genes enriched based on multiple pathways. Pathway enrichment led to the prediction of genes involved in the DDR mechanism interacting with the other query genes present in multiple DDR pathways based on WikiPathway 2021 Human, Reactome 2022, PFOCR pathways, and KEGG 2021 Human pathways, as shown in the bar chart (see **Fig. 4 A**). The query genes associated with multiple enriched DDR pathways can be found in **Supplementary Table 2** and are arranged in ascending order based on the combined score. Enrichr KG provides pathway association data based on p values, q-values, z scores, and combined scores to prioritize the genes involved in specific pathways (John et al. 2023).

**Figure 4:**
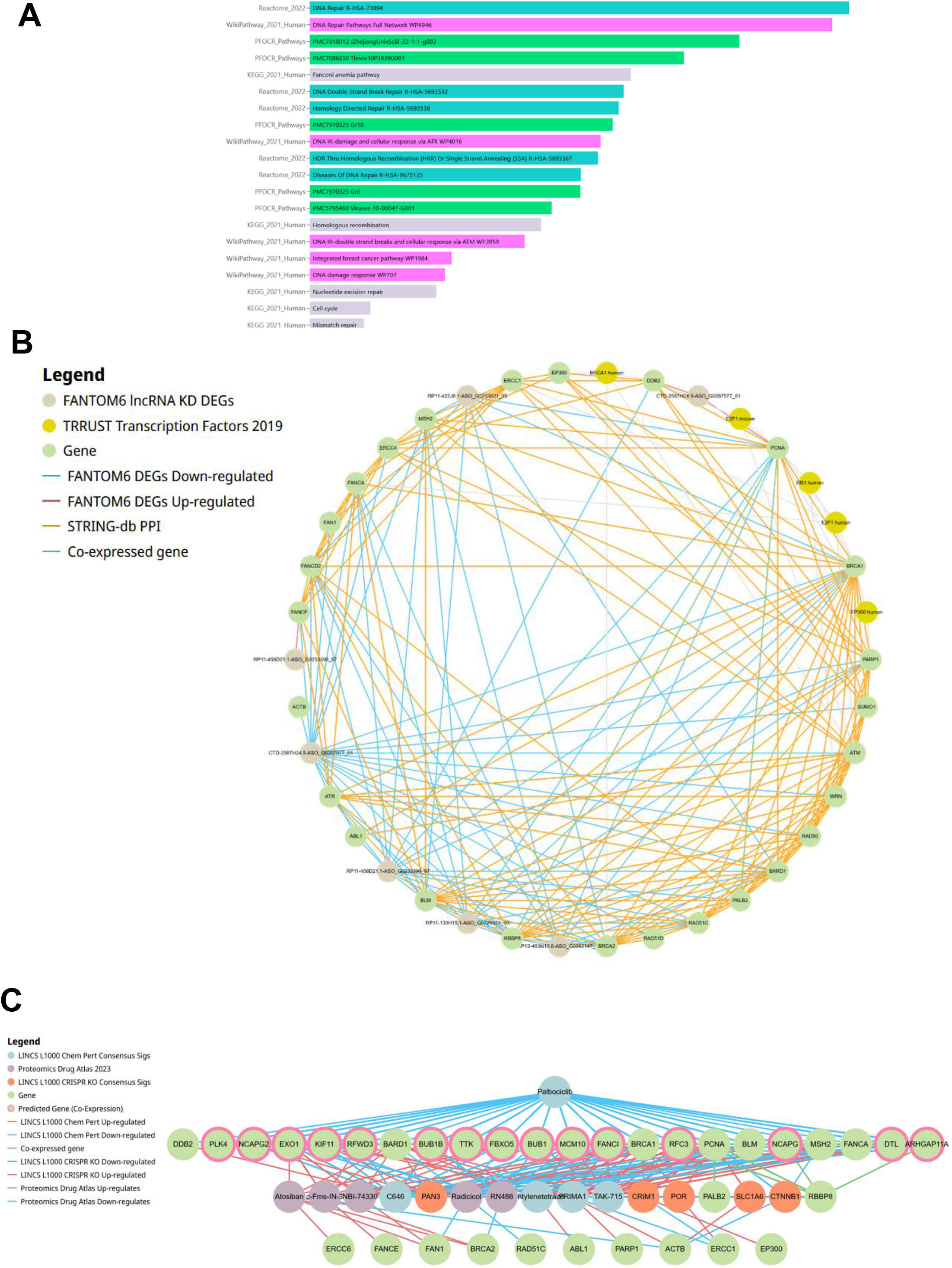
**MiR-interacting genes and gene set enrichment analysis (miRNet). A**) Bar chart representation of gene enrichment leading to participation of query genes in multiple pathway databases; **B**) Circular representation of genes enriched for KD DEGs, transcription factors, downregulated and upregulated DEGs, protein‒protein interactions, and coexpressed genes based on enrichment of multiple databases; **C**) Chem_Compounds-genes interactions leading up/downregulation of genes predicted by LINCs L 1000; blue lines correspond to downregulation, and orange lines indicate upregulation of genes by the interactions.

Further enrichment of the genes based on the FANTOM6 subnetwork inferred the genes associated with functional elements, including transcription and DEG mechanisms. This was obtained by annotating long noncoding RNAs (lncRNAs) to predict upregulated and downregulated DEGs. The blue lines in **Fig. 4B** linking the genes refer to downregulated DEGs, red lines to upregulated DEGs (PCNA, DDB2, FANCF interacting with KD), and gray circles indicate successful knockdown (KD) DEGs. The green lines connecting PCNA with PALB2 and RBBP8 with KD lncRNA indicate the coexpressed genes. The orange lines in **Fig. 4B** specify protein‒protein interactions between the linked genes. We found that the BRCA1 and ATM genes had the highest number of protein‒protein interactions based on the network.

The results shown in **Fig. 4C** refer to palbociclib (blue circle) downregulating multiple coexpressed genes by direct interaction (shown as blue lines). Orange circles indicate consensus knockouts (KOs) based on LINCS L 1000 CRISPR KO consensus Sigs prediction. The orange lines in **Fig. 4C** indicate the chemical compound (purple circles) interactions leading to the upregulation of genes as obtained by reference to the Proteomics Drug Atlas.

### 3.7. Ligands

Among the 330 ligands searched from the PSRR web server (https://rnadrug.shinyapps.io/PSRR/), with a score of >0.85. The compounds were subjected to SWISSADME analysis to filter based on multiple druggable parameters, resulting in 113 compounds. The obtained structures were then docked to be filtered based on their binding energy scores. Forty-two compounds showed possible interactions with binding energies >-5 kcal/mol. The top 13 compounds interacting with prioritized miRs are shown in **Table 4** and **Table 5**. (Kim et al., 2021)

**Table 4:** Ligands targeting mRNA-miR duplexes obtained from PSRR web server and PubChem.

| Ligand No. | Ligand Name | Ligand CID |
| --- | --- | --- |
| 1 | Cyclonite | 8490 |
| 2 | Reversine | 210332 |
| 3 | Trichostatin A | 444732 |
| 4 | Dexamethasone | 5743 |
| 5 | Betamethasone | 9782 |
| 6 | Diflorasone | 71415 |
| 7 | Fluorouracil | 3385 |
| 8 | D-Galactose | 6036 |
| 9 | Gemcitabine | 60750 |
| 10 | Floxuridine | 5790 |
| 11 | Atorvastatin | 60823 |
| 12 | (3R,5S)-Atorvastatin | 46780495 |
| 13 | (3S,5R)-Atorvastatin | 6093359 |

**Table 5:**
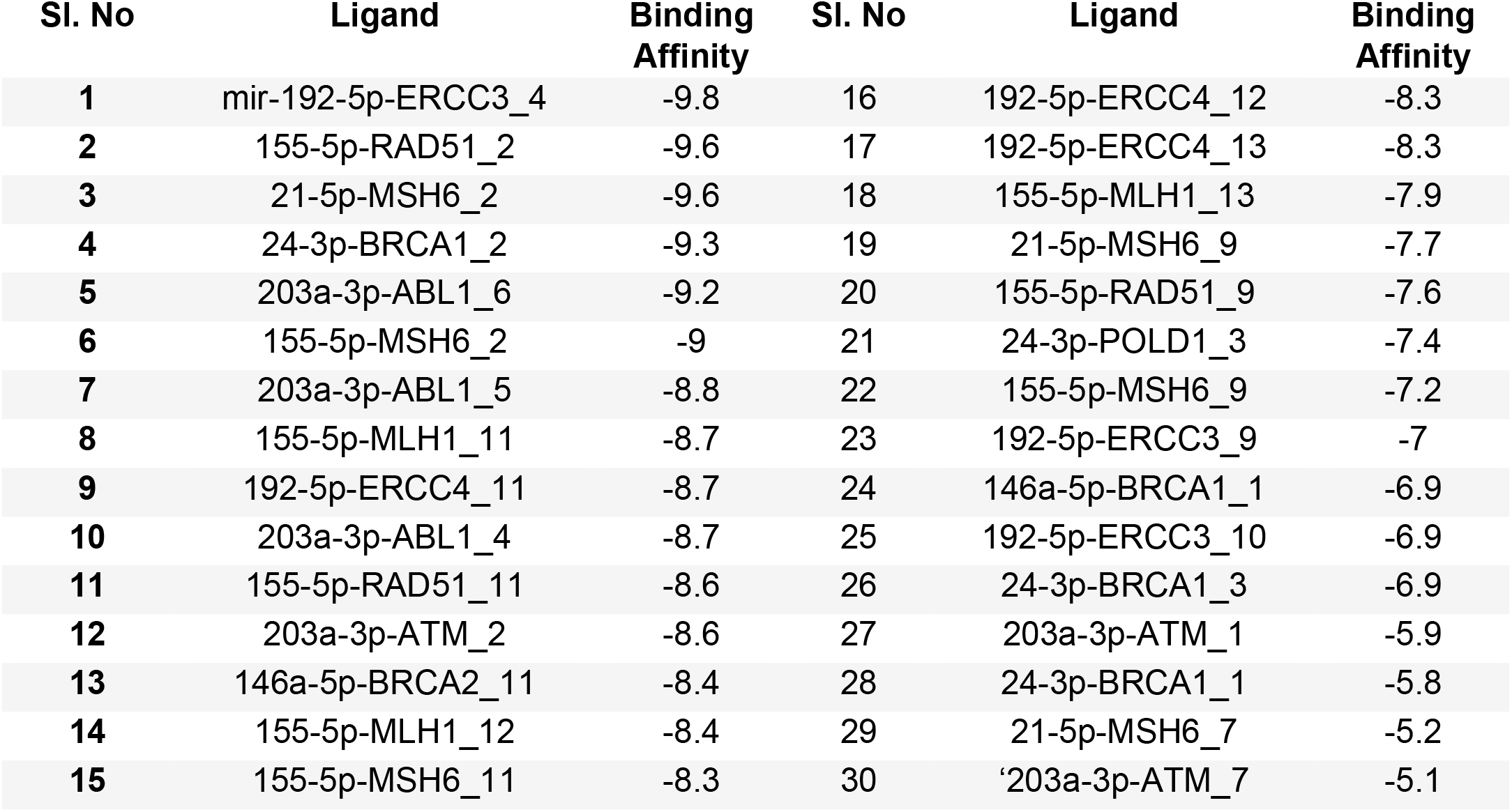
Docking scores (binding affinity) obtained from AutoDock Vina. (Ligand Notation: miRNA_mRNA_Ligand No.)

### 3.7. Molecular Docking and Analysis

Docking studies between the mRNA-miR duplexes and target ligands were conducted using AutoDock Vina. Since the binding site was not known, the grid box was adjusted to cover the surface of the entire nucleic acid. Out of all the binding poses in each simulation, the pose with the highest (most negative) binding energy was selected. These results are shown in **Table 5**. The binding poses of interactions having a binding energy > -8.5 kcal/mol visualized on PyMOL are shown in **Fig. 5 (**see also **Table 5)**. The list of hydrogen bonds between nucleotides and ligands with a distance of less than 3 Å was generated by Discovery Studio (**Suppl. Table 3**). These bonds indicate good binding with the nucleotides. Binding energies of > -8.5 kcal/mol indicate strong binding and high affinity. This indicates the inhibitory action of the mRNA/gene interaction under consideration. It has been established that ligand-miR complexes with higher binding scores are comparatively more highly stabilized than complexes with lower binding scores. These predicted compounds are expected to have downregulating effects, and because of their high binding score, they would indicate a higher downregulation of the miRs.

**Figure 5:**
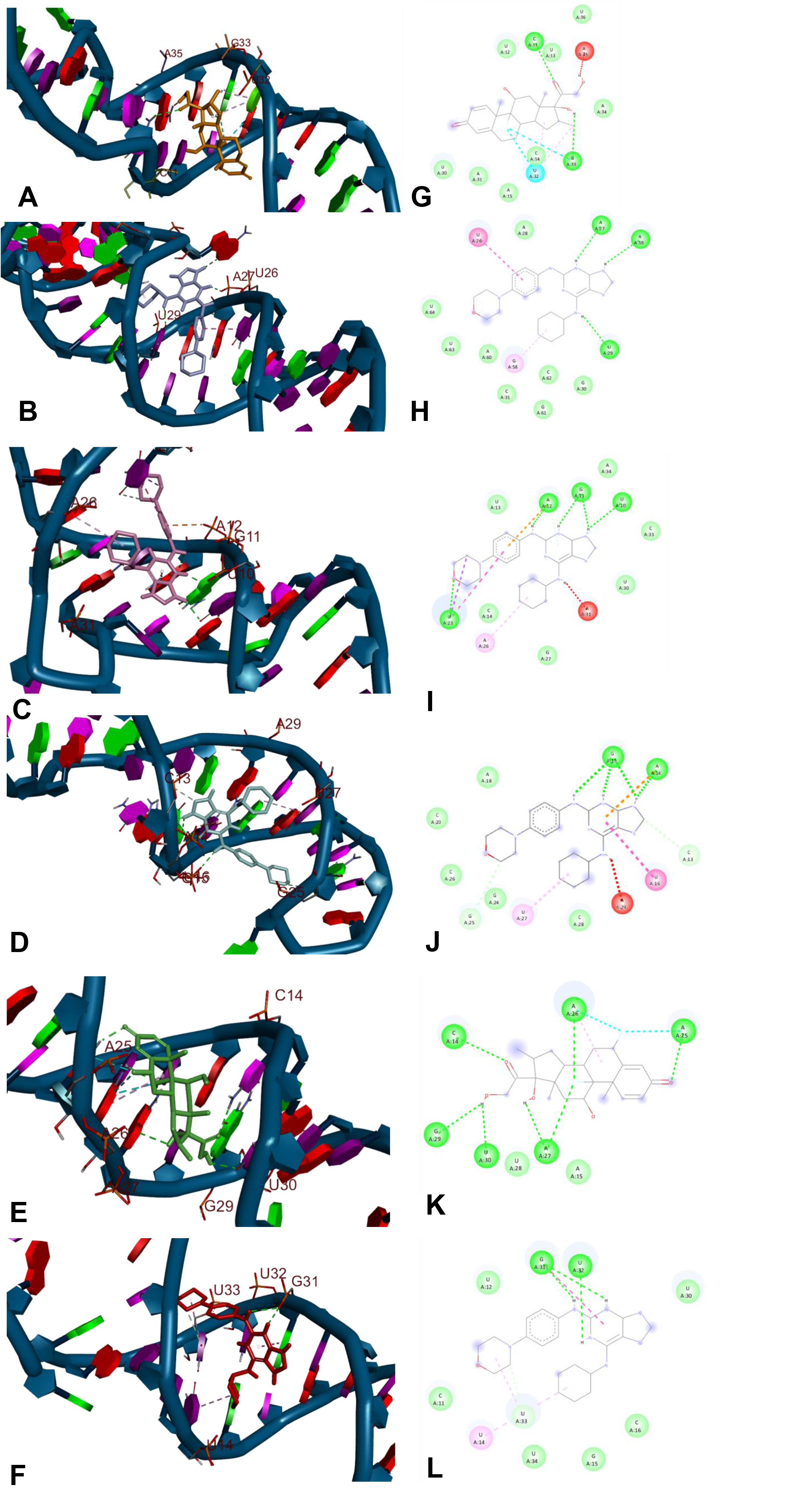
**Schematic representation of interacting ligands with the miRNA‒ mRNA duplex with the best binding score. A**) 192-5p-ERCC3 ligand 4, **B**) 155-5p- RAD51 ligand 2, **C**) 21-5p-MSH6 ligand 2, **D**) 24-3p-BRCA1 ligand 2, **E**) 203a-3p- ABL1 ligand 6, and **F**) 203a-3p-ATM_2. **G, H, I, J, K, and L** show 2D interactions of the ligands with binding site residues.

## 4. DISCUSSION

Oncogenesis, metastasis, and resistance to particular therapies have all been linked to miRs (Peng et al., 2013; Rani et al., 2013; van Schooneveld et al., 2015). miRs have been found to inhibit the translation of RNAs encoding proteins involved in DDS and DDR mechanisms; thus, more verified miR targets have to be discovered. Herein, we found drugs with the best DDR-associated miR binding properties. These include kinase inhibitors, glucocorticoid steroids, and statins. These compounds are already predicted to act as downregulators of miRs and therefore could serve as structural platforms for the design of novel cancer therapeutics.

The compounds found in our study are all drugs that have already been approved by regulatory agencies for other purposes and could thereby be repurposed for use in the treatment of several cancers and deserve validation by further *in vitro* and *in vivo* studies. For instance, downregulation of the BRCA1 and BRCA2 genes leads to higher susceptibility to breast cancer (Venkitaraman, 2002). ERCC3 and ERCC4 are essential genes for NER, and their downregulation has been shown to lead to hepatocellular carcinoma (Xie et al., 2011). Downregulation of ATM is involved in various cancers, including nasopharyngeal carcinoma (Bose et al., 2009). MLH1 and MSH6 downregulation has been linked to colorectal cancer (Edwards et al., 2009; Hemminki et al., 1994), and RAD51 is downregulated in cancers and increases sensitivity to tumors in the case of hypoxia (Bindra et al., 2004). Moreover, POLD1 downregulation has been linked to cognitive function impairment in cases of Alzheimer’s disease (Gao et al., 2019; Song et al., 2015). The miRs found in this study have been shown to downregulate these genes, which in many cases would lead to cancer development.

Furthermore, PALB2 is involved in homologous recombination repair of double-strand breaks, plays a crucial role as a tumor suppressor, and maintains genome integrity. Mutations in PALB2 increase the risk of breast, pancreatic, and ovarian cancers (Nepomuceno et al., 2017). PCNA is essential in DNA replication and NER mechanisms by repairing DNA damage caused by exposure to UV light and carcinogens (Essers et al., 2005). Downregulation/mutations in PCNA are highly correlated with colorectal and breast cancer occurrence. RBBP8 has a fundamental role in DNA replication, transcription, and DDR mechanisms (Mijnes et al., 2018). These three genes (PALB2, PCNA, and RBBP8) are closely associated with BRCA1 and BRCA2, leading to breast, colorectal, and endometrial cancers.

Mechanistically, ATM is a gene encoding ATM serine/threonine kinase, a DDS protein kinase that is activated in the presence of DNA strand breaks and hence is essential to trigger the DDR process. miR-421 has been shown to regulate ATM by binding to the 3’-UTR of ATM-encoding mRNA, which has been linked to increased cell radiosensitivity (Hu et al., 2010). Further research has revealed that tumors such as neuroblastomas are accompanied by an upregulation of miR-421 or overexpression of miR-24, which targets and downregulates histone H2AX expression, resulting in an inefficient DDR process (Hu et al., 2010; Lal et al., 2009). Consequently, to prevent carcinogenesis, it is necessary to find drugs that inhibit the regulatory action of miRs on DDS and DDR protein expression.

The identified genes and miRs were queried using multiple platforms to predict and annotate their involvement in interfering with DDS/DDR processes. NDEx IQuery augmented the specific genes involved in the integrated networks by ranking the genes based on multiple scores into multiple pathways, including intrinsic apoptotic signaling pathway response to DNA damage, DNA damage checkpoints, mismatch repair, double-strand break repair (DSBR), DSBR via homologous recombination, interstrand cross-link repair, mitotic DNA integrity checkpoint, NER, DNA incision, postreplication repair, telomere maintenance, and regulation of DDR. This is expected to lead to an understanding of the involvement of these genes in specific pathways and the interacting genes involved in specific DDS and DDR mechanisms.

Network analysis of query genes using miRNet leveraged miR-disease associations by integrating the gene and disease information stored in multiple databases and by incorporating transcription factor and single nucleotide polymorphism data affecting the function of miRs. It facilitated the ranked miR visualization, exhibiting their association with diseases based on information in the DisGeNET database and multiple scoring. The range of disorders, diseases, and cancer types was integrated into the disease-miR network (**see Suppl. Fig. 1D**).

Gene enrichment analysis is an important step in analyzing multiomics-based experimental data sets. This enriched data facilitated the understanding of the specific role of the genes involved in DDR mechanisms, as it collates the information linked to the query genes from multiple databases and helps to visualize them in an infographic (see **Fig. 4A)**. It also highlights the gene‒gene interactions, coexpressed genes, upregulated/downregulated genes and transcriptome-associated DEGs (**see Fig. 4B**). Furthermore, as shown in **Fig. 4C**, it augments the upregulated/downregulated genes due to their interaction with specific chemical compounds based on color-coded lines. This piece of information may be of specific interest to the researchers involved in drug design and development.

The associations between the genes, pathways, drugs, and diseases can be oriented based on the collating transcriptomics data using the Library of Integrated Network-Based Cellular Signatures (LINCS) portal. It provides information on assays, cell types, and perturbagens based on varied links leading to the upregulation and downregulation of genes. The green circles with pink-bordered circles indicate coexpressed genes. Knowledge of the molecular mechanism by which small ligands can act as perturbagens is crucial to understanding the cellular response to drugs. LINCS L1000 builds a network based on comprehensive information on gene expression and cellular processes exposed to a wide range of perturbagens. It provides information on the upregulation or downregulation of the genes as an effect of the perturbagen interaction. LINCS shows that the PAN3 (Up), CRIM1 (Up), POR (Up), SLC1A6 (Up), and CTNNB1 (Down) genes are up/downregulated due to perturbagens. For example, our data (**Fig 4C)** show that downregulation of CTNNB1 leads to inhibition of the Wnt/β-catenin signaling pathway by downregulating the Axin2, LEF1, and CCND1 genes. In agreement with these data, Zhou et al. reported that CTNNB1 knockdown leads to inhibition of H295R cell proliferation and a decline in aldosterone secretion as a response of H295R cells to Ang II by inhibiting the Wnt/β-catenin signaling pathway. This indicates the importance of the Wnt/β- catenin signaling pathway as a crucial target to reduce aldosterone secretion in the therapeutics of aldosterone-producing adenomas (Zhou et al., 2020). A similar enrichment using the Proteomics Drug Atlas 2023 (PDA-2023) prediction tool showed the associations of the compounds involved in the downregulation of certain genes. For instance, palbociclib downregulates several genes, including BARD1, PCNA, BRCA1, MSH2, DDB2, FANCA, and BLM, as illustrated by the blue line in **Fig 4C**. In addition, atosiban upregulates genes such as ERCC6, FAN1, BRCA2, and BARD1. Integrated database network-based predictions are gaining significance in the discovery of novel therapeutics for diseases and cancers (Mitchell et al., 2023). PDA- 2023 has made a significant effort in collating proteome-wide effects in the form of fingerprints belonging to 875 chemical compounds as perturbagens as a knowledge base to understand the mechanism of action and drug repurposing.

Computational drug discovery (CDD) can reduce the time taken in the research cycle and reduce the costs involved. Novel discovery of drugs and the development process can cost up to a billion dollars (Myers & Baker, 2001) and can be costly and time-consuming. Structure-based drug design (SBDD), ligand-based drug design (LBDD), and sequence-based techniques are the three most common CDD approaches. There have been steadfast developments in CDD and high-throughput screening, allowing access to gigantic libraries of compounds (ligands) to be screened and synthesized in a short period (Tang et al., 2006), (Lahana, 1999; Lobanov, 2004). CDD is the term used to represent the tools and libraries used to store, manage, analyze, and model compound-target (protein/RNA) interactions. The use of these tools has generally reduced costs by approximately 50% (Tan et al., 2010).

As a limitation of our study, we can mention that the docking methods used to study protein‒ligand interactions are thereby being modified (or new programs are being created altogether) to model RNA-ligand interactions (Wehler & Brenk, 2017). However, when transferring the procedures, there are several obstacles to consider. For example, RNA molecules are strongly charged, resulting in strong ionic solvation and interaction (N. Foloppe & Hubbard, 2006; Hermann, 2002; Thomas & Hergenrother, 2008). At the binding site, water molecules and ions are frequently detected. During docking, these must be considered because they may either mediate or be displaced by the ligand. Thus, ligand binding might result in conformational changes or induced fit movements, which must be taken into account when modeling RNA–ligand interactions (Hermann, 2002).

Finally, depending on the selected descriptors and type specification, scoring functions that were parameterized on protein‒ligand complexes (*i.e.,* empirical and knowledge-based scoring functions) must be reparameterized on RNA-ligand complexes (Nicolas Foloppe et al., 2006). Considering these difficulties, researchers have chosen to either adapt docking systems designed for protein‒ligand docking to RNA or build new methodologies and scoring functions. In one of the first applications of RNA-ligand SBVS in the screening of ligands that prevented the formation of the transactivation response (TAR) element RNA-Tat complex, Filikov et al. used modified versions of DOCK and ICM (Filikov et al., 2000). In that study, the complex was disrupted by three different compounds. In a later study, evaluation and docking were improved (Lind et al., 2002). Remarkably, 11 ligands that bind to TAR were discovered in this study; for some of them, cell activity could be shown, and nuclear magnetic resonance was used to confirm binding to the intended binding site (Lind et al., 2002).

## 5. CONCLUDING REMARKS

Our computational study showed the correlation between several DDS and DDR genes and the effect of their downregulation on cell transformation. We also studied the effect of gene silencing by miRs and their consequences, constructed the structure of various miR-mRNA complexes, and used these structures to study the binding and effects of ligands possibly downregulating these complexes. This binding reduces the effect of gene silencing in essential genes for DDS and DDR processes. Our gene enrichment analysis was effective in identifying the associations of genes and miR associations in DDS and DDR signaling pathways, miR-disease interactions, gene‒ gene interactions, and gene‒ligand associations (leading to up/downregulation). The predicted miRs may also serve as significant prognostic biomarkers and contribute to future therapeutics. Interestingly, the 13 identified ligands through our docking studies belong to classes of drugs that are approved as therapeutics. Data gathered from this study will serve as a starting point for the repurposing of old drugs as novel disease/cancer therapeutics.

## NOTES

**Funding:** This study was funded by the National Fund for Scientific and Technical Investigations (FONCYT), Argentina [PICT-2021-I-A-0147] to DCR.

**The Role of the Funder:** The funders had no role in the design of the study; the collection, analysis, or interpretation of the data; the writing of the manuscript; or the decision to submit the manuscript for publication.

**Author Disclosures:** RSB, VS, LBS, RDC and GMSE have no conflicts of interest to disclose.

**Author contributions:** RSB, VS, and LBS performed computational drug discovery analysis and data analysis. RDC and GMSE participated in the discussion and manuscript preparation. All the authors edited the manuscript and discussed and agreed on its content.

**Disclaimers:** not applicable

### Any Prior presentations: **none**

**Data Availability:** The authors declare that all the relevant data supporting the findings of this study are available within the paper and its supplementary information and from the corresponding authors upon reasonable request. The code is available from the corresponding authors upon reasonable request.

## Acknowledgments

The authors greatly acknowledge the management of Manipal Institute of Technology, Manipal; Manipal Academy of Higher Education, Manipal, India; Carnegie Mellon University, PES University, Pennsylvania; and the CONICET- National University of San Luis for their valuable support for this work.

## SUPPLEMENTARY FIGURE LEGENDS

**Supplementary Figure 1: Schematic representation of the miR-gene-disease interaction networks. A**) miR-gene interaction network of target genes, **B**) miRs with highest betweenness scores oriented, **C**) augmented miR interaction with genes, and **D**) miRNA-disease associations based on DisGeNET database data and ranked by scores.

**Supplementary Figure 2: Secondary structure generated using the RNAfold Web Server of miRNA‒mRNA duplexes. A**) BRCA1-24-3p, **B**) POLD1-24-3p, **C**) ABL1-203a-3p, **D**) ATM-203a-3p, **E**) MSH6-21-5p, **F**) ERCC3-192-5p, **G**) ERCC4- 192-5p, **H**) BRCA1-146a-5p, **I**) BRCA2-146a-5p, **J**) MLH1-155-5p, **K**) MSH6-155-5p, and **L**) RAD51-155-5p.

