## Supplementary figures and images for "Drugs targeting microRNAs that regulate DNA damage sensing and repair mechanisms: *A computational drug discovery approach*"

### SUPPL. FIGURES

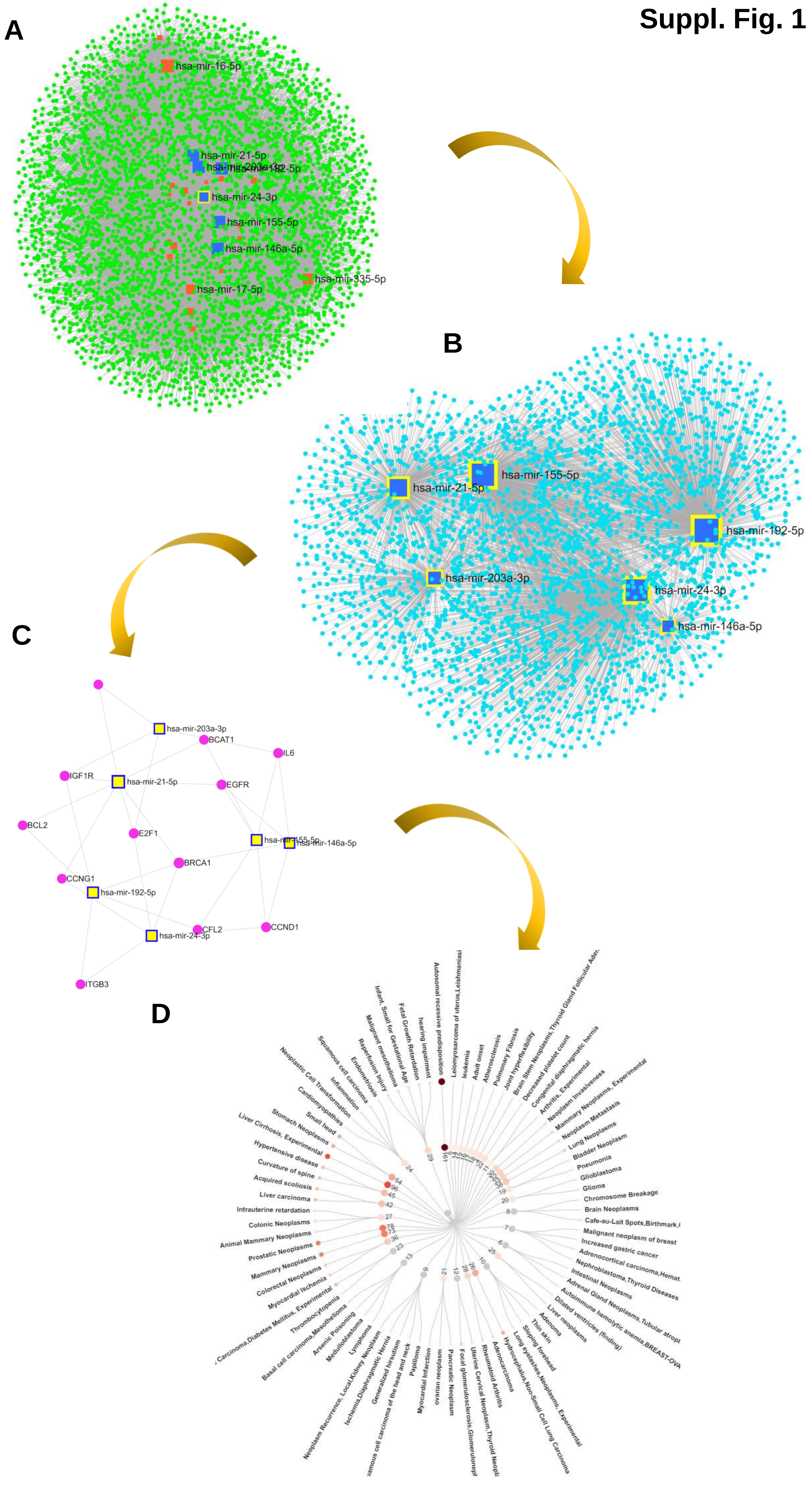

**Suppl. Figure 2**

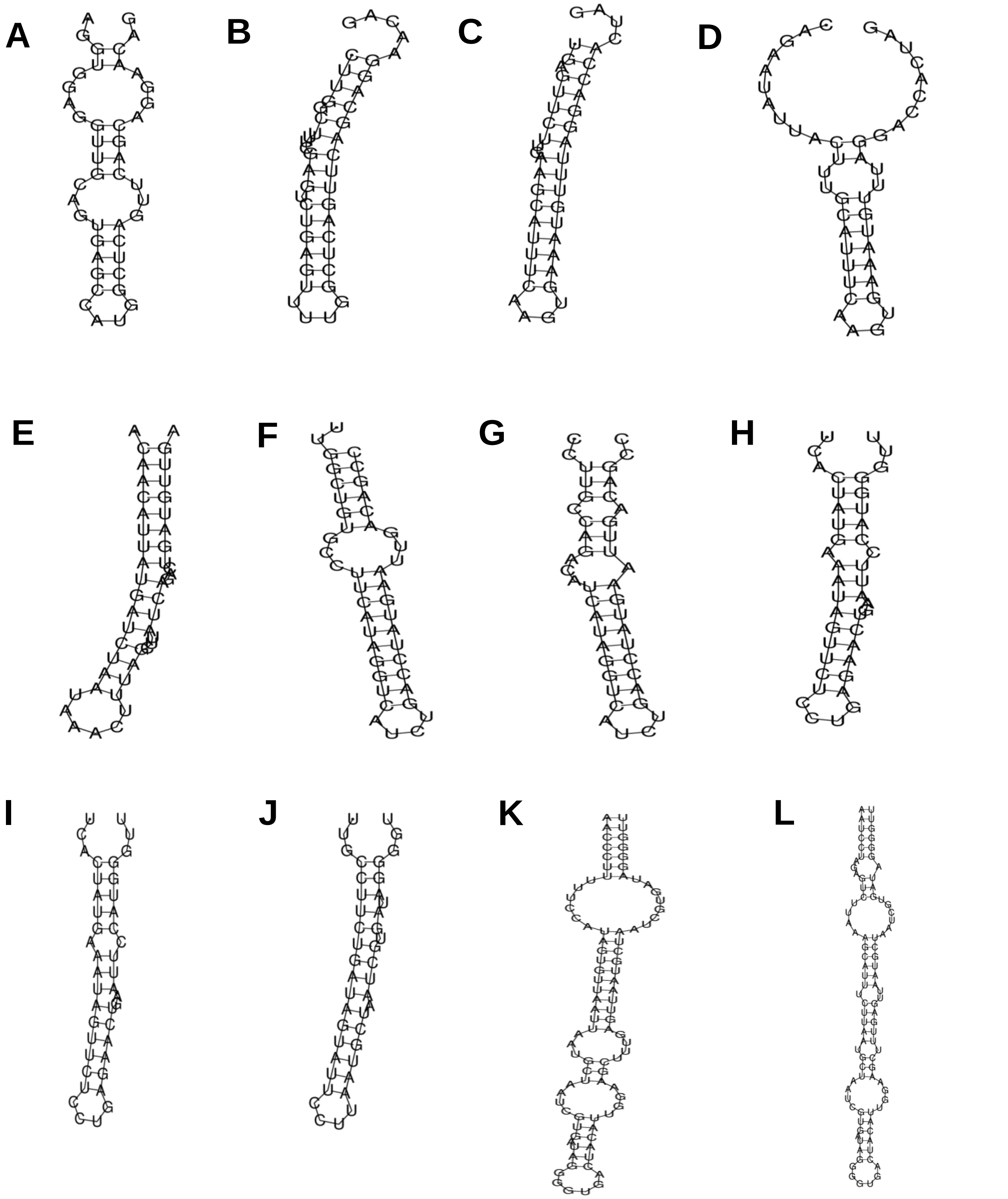
