## Supplementary material for "Drugs targeting microRNAs that regulate DNA damage sensing and repair mechanisms: *A computational drug discovery approach*": SUPPL. TABLES

**Supplementary Table 1:** Prioritized miRs and their involvement in diseases

| **Disease (DisGeNET)** | **Hits** | **Pval** |
|---|---|---|
| Reperfusion Injury | 29 | 1.33E-10 |
| Myocardial Ischemia | 36 | 7.45E-09 |
| Neoplasm Invasiveness | 30 | 9.61E-09 |
| Mammary Neoplasms | 73 | 2.65E-08 |
| Prostatic Neoplasms | 78 | 3.12E-08 |
| Mammary Neoplasms, Experimental | 32 | 3.79E-08 |
| Stomach Neoplasms | 54 | 0.000000336 |
| Malignant mesothelioma | 29 | 0.000000718 |
| Liver carcinoma | 42 | 0.000000864 |
| Neoplasm Metastasis | 33 | 0.00000139 |
| Glioblastoma | 20 | 0.00000183 |
| Hydrocephalus | 26 | 0.00000206 |
| Non-Small Cell Lung Carcinoma | 26 | 0.00000206 |
| Cardiomyopathies | 24 | 0.00000217 |
| Adenocarcinoma | 28 | 0.00000259 |
| Uterine Cervical Neoplasm | 12 | 0.00000305 |
| Thyroid Neoplasm | 12 | 0.00000305 |
| Animal Mammary Neoplasms | 27 | 0.0000033 |
| Adenoid Cystic Carcinoma | 23 | 0.00000397 |
| Diabetes Mellitus, Experimental | 23 | 0.00000397 |
| Neoplastic Cell Transformation | 24 | 0.00000467 |
| Pancreatic Neoplasm | 21 | 0.0000133 |
| Papilloma | 9 | 0.0000133 |
| Dilated ventricles (finding) | 25 | 0.0000201 |
| leukemia | 14 | 0.0000278 |
| Small head | 54 | 0.0000361 |
| Inflammation | 24 | 0.0000478 |
| Lung Neoplasms | 38 | 0.0000492 |
| Decreased platelet count | 22 | 0.0000591 |
| Glioma | 20 | 0.0000706 |
| Curvature of spine | 45 | 0.0000708 |
| Atherosclerosis | 16 | 0.0000841 |
| Colorectal Neoplasms | 36 | 0.0000842 |
| Squamous cell carcinoma | 24 | 0.000085 |
| Liver neoplasms | 25 | 0.0000855 |
| Lymphoma | 13 | 0.0000885 |
| Malignant neoplasm of breast | 7 | 0.0000936 |
| Infant, Small for Gestational Age | 29 | 0.000101 |
| Adult onset | 15 | 0.000102 |
| Colonic Neoplasms | 27 | 0.000108 |
| Adrenal Gland Neoplasms | 6 | 0.000129 |
| Tubular atrophy | 6 | 0.000129 |
| Transitional cell carcinoma of bladder | 6 | 0.000129 |
| ovarian neoplasm | 21 | 0.000141 |

| | | |
|---|---|---|
| Liver Cirrhosis, Experimental | 96 | 0.000194 |
| Bladder Neoplasm | 19 | 0.000238 |
| Increased gastric cancer | 7 | 0.000267 |
| Chromosome Breakage | 8 | 0.000343 |
| Hypertensive disease | 45 | 0.000347 |
| Adenoma | 10 | 0.000348 |
| Basal cell carcinoma | 13 | 0.000353 |
| Mesothelioma | 13 | 0.000353 |
| Squamous cell carcinoma of the head and neck | 9 | 0.000364 |
| Fetal Growth Retardation | 29 | 0.000371 |
| Focal glomerulosclerosis | 12 | 0.000429 |
| Glomerulonephritis | 12 | 0.000429 |
| Infarction, Middle Cerebral Artery | 12 | 0.000429 |
| Intestinal Neoplasms | 6 | 0.000441 |
| Acquired scoliosis | 42 | 0.000468 |
| Congenital diaphragmatic hernia | 11 | 0.000511 |
| Thrombocytopenia | 23 | 0.000519 |
| Intrauterine retardation | 27 | 0.00052 |
| Medulloblastoma | 13 | 0.000529 |
| Myocardial Infarction | 21 | 0.000553 |
| Thin skin | 10 | 0.000593 |
| Adrenocortical carcinoma | 7 | 0.000628 |
| Hematologic Neoplasms | 7 | 0.000628 |
| Ischemia | 9 | 0.000659 |
| Diaphragmatic Hernia | 9 | 0.000659 |
| Leiomyosarcoma of uterus | 5 | 0.000666 |
| Leishmaniasis, Cutaneous | 5 | 0.000666 |
| Brain Neoplasms | 8 | 0.000685 |
| Brain Stem Neoplasms | 4 | 0.000775 |
| Thyroid Gland Follicular Adenoma | 4 | 0.000775 |
| Growth hormone excess | 4 | 0.000775 |
| Pituitary growth hormone cell adenoma | 4 | 0.000775 |
| Shared Paranoid Disorder | 4 | 0.000775 |
| Arthritis, Experimental | 11 | 0.000803 |
| Long eyelashes | 10 | 0.000964 |
| Neoplasms, Experimental | 10 | 0.000964 |
| Rheumatoid Arthritis | 28 | 0.00105 |
| Arsenic Poisoning | 13 | 0.00111 |
| Neoplasm Recurrence, Local | 9 | 0.00112 |
| Kidney Neoplasm | 9 | 0.00112 |
| Autoimmune hemolytic anemia | 6 | 0.00114 |
| Breast-Ovarian cancer | 6 | 0.00114 |
| Pleural Diseases | 6 | 0.00114 |
| Hearing impairment | 29 | 0.00115 |
| Cafe-au-Lait Spots | 8 | 0.00125 |
| Birthmark | 8 | 0.00125 |
| Colitis | 8 | 0.00125 |
| Nephroblastoma | 7 | 0.00129 |

| | | |
|---|---|---|
| Thyroid Diseases | 7 | 0.00129 |
| Endometriosis | 24 | 0.00143 |
| Sloping forehead | 10 | 0.00151 |
| Pneumonia | 19 | 0.00156 |
| Joint hyperflexibility | 18 | 0.00162 |
| Autosomal recessive predisposition | 161 | 0.00165 |
| Pulmonary Fibrosis | 17 | 0.00166 |
| Generalized hirsutism | 9 | 0.00182 |

**Supplementary Table 2**: Enrichment of genes associated with multiple DDR pathways

| Term | Library | p-value | q-value | z-score | combined score |
|---|---|---|---|---|---|
| **DNA Repair R-HSA-73894** | Reactome_2022 | 6.40E-52 | 1.68E-49 | 703.1 | 82880 |
| **DNA Repair Pathways Full Network WP4946** | WikiPathway_2021_Human | 2.61E-50 | 2.58E-48 | 653.7 | 74630 |
| **PMC5795460 Viruses-10-00047-G001** | PFOCR_Pathways | 1.14E-23 | 3.75E-21 | 788.8 | 41670 |
| **PMC7818012 JZhejiangUnivSciB-22-1-1-g002** | PFOCR_Pathways | 1.70E-41 | 2.78E-38 | 440.6 | 41360 |
| **PMC7086350 Thnov10P3939G001** | PFOCR_Pathways | 3.06E-36 | 2.51E-33 | 411.9 | 33680 |
| **Fanconi anemia pathway** | KEGG_2021_Human | 3.48E-31 | 3.02E-29 | 425.8 | 29860 |
| **Diseases Of DNA Repair R-HSA-9675135** | Reactome_2022 | 2.01E-26 | 1.06E-24 | 340.9 | 20170 |
| **Homologous recombination** | KEGG_2021_Human | 1.16E-22 | 5.04E-21 | 332.3 | 16780 |
| **DNA IR-damage and cellular response via ATR WP4016** | WikiPathway_2021_Human | 2.61E-28 | 1.29E-26 | 255.2 | 16210 |
| **Homology Directed Repair R-HSA-5693538** | Reactome_2022 | 4.80E-30 | 4.21E-28 | 206.9 | 13970 |
| **PMC7970325 Gr10** | PFOCR_Pathways | 1.80E-29 | 9.83E-27 | 190.1 | 12580 |
| **DNA Double-Strand Break Repair R-HSA-5693532** | Reactome_2022 | 1.63E-30 | 2.14E-28 | 181.7 | 12460 |
| **HDR Thru Homologous Recombination (HRR)** | Reactome_2022 | 4.46E-28 | 2.93E-26 | 190.8 | 12020 |
| **PMC7970325 Gr8** | PFOCR_Pathways | 2.14E-26 | 8.78E-24 | 184 | 10880 |
| **DNA IR-double strand breaks and cellular response via ATM WP3959** | WikiPathway_2021_Human | 4.32E-21 | 1.43E-19 | 226.4 | 10620 |
| **Nucleotide excision repair** | KEGG_2021_Human | 1.01E-12 | 2.93E-11 | 134.1 | 3704 |
| **DNA damage response WP707** | WikiPathway_2021_Human | 1.51E-13 | 3.00E-12 | 106.2 | 3134 |

| | | | | | |
|---|---|---|---|---|---|
| **Integrated breast cancer pathway WP1984** | WikiPathway_2021_Human | 3.80E-14 | 9.39E-13 | 60.7 | 1876 |
| **Mismatch repair** | KEGG_2021_Human | 7.086E-06 | 0.000123 | 99.73 | 1183 |
| **Cell cycle** | KEGG_2021_Human | 1.745E-06 | 3.79E-05 | 29.78 | 394.9 |

**Supplementary Table 3:** Interacting bonds and bond lengths between nucleotides and ligands. (*)

| ***Bond Name*** | ***Bond Length (Å)*** | ***Bond Name*** | ***Bond Length (Å)*** |
|---|---|---|---|
| **21-5p-MSH6-Ligand 2** | | **155-5p-MSH6-Ligand 11** | |
| A:U23:H3 - :UNL1:O | 2.76756 | A:C3:H42 - :UNL1:O | 2.88433 |
| :UNL1:HN - A:G11:N7 | 1.86413 | :UNL1:H - A:G87:O6 | 1.97436 |
| :UNL1:HN - A:G11:O6 | 2.81238 | :UNL1:H - :UNL1:O | 2.01747 |
| :UNL1:HN - A:U10:O4 | 2.38618 | **155-5p-RAD51-Ligand 2** | |
| **21-5p-MSH6-Ligand 7** | | :UNL1:HN - A:A27:OP2 | 2.38734 |
| A:G35:H21 - :UNL1:O | 2.02448 | :UNL1:HN - A:A59:N3 | 2.48991 |
| A:A40:H61 - :UNL1:O | 1.86905 | :UNL1:H - A:U29:O4 | 2.43053 |
| :UNL1:H - A:G39:O6 | 2.88985 | **155-5p-RAD51-Ligand 9** | |
| **21-5p-MSH6-Ligand 9** | | :UNL1:HN - A:A59:N3 | 2.85609 |
| A:A6:H61 - :UNL1:O | 2.3033 | :UNL1:HN - A:A27:OP2 | 2.41842 |
| A:C37:H41 - :UNL1:O | 2.81743 | :UNL1:H - A:A27:OP2 | 2.3144 |
| :UNL1:HN - A:U41:O4 | 2.65038 | :UNL1:H - A:G61:O6 | 2.842 |
| :UNL1:H - A:A40:N7 | 2.13489 | **155-5p-RAD51-Ligand 11** | |
| :UNL1:H - A:U38:O4 | 2.13828 | A:A28:H62 - :UNL1:O | 2.52324 |
| :UNL1:H - A:U8:O4 | 2.2019 | A:A27:H61 - :UNL1:O | 2.5175 |
| **24-3p-BRCA1-Ligand 1** | | A:A27:H62 - :UNL1:O | 2.72984 |
| :UNL1:O - A:U27:O4 | 2.37816 | :UNL1:H - A:U29:O4 | 2.51039 |
| :UNL1:O - A:U16:O4 | 2.39976 | :UNL1:H - :UNL1:O | 2.8712 |
| **24-3p-BRCA1-Ligand 2** | | **192-5p-ERCC3-Ligand 4** | |
| :UNL1:HN - A:G15:N7 | 2.31018 | A:C11:H41 - :UNL1:O | 2.70012 |
| :UNL1:HN - A:G15:N7 | 2.51759 | :UNL1:H - A:G33:O6 | 2.50444 |
| :UNL1:HN - A:A14:N7 | 2.3678 | **192-5p-ERCC3-Ligand 9** | |
| :UNL1:H - A:G15:OP2 | 2.48671 | A:C28:H41 - :UNL1:O | 2.3642 |
| **24-3p-BRCA1-Ligand 3** | | A:C28:H42 - :UNL1:O | 2.30035 |
| A:G5:H1 - :UNL1:O | 2.45392 | A:C29:H41 - :UNL1:O | 2.19108 |
| A:A41:H61 - :UNL1:O | 2.43284 | :UNL1:H - A:G26:O6 | 2.45038 |
| A:A41:H62 - :UNL1:O | 2.72442 | :UNL1:H - A:A17:N7 | 2.04012 |
| :UNL1:H - A:G6:O6 | 2.917 | **192-5p-ERCC3-Ligand 10** | |
| :UNL1:H - A:G6:O6 | 2.0755 | A:A31:H61 - :UNL1:O | 2.31813 |
| :UNL1:H - A:G5:O6 | 2.12818 | A:A34:H62 - :UNL1:O4* | 2.76614 |
| **24-3p-POLD1-Ligand 3** | | A:C14:H41 - :UNL1:O | 2.43213 |
| A:C36:H41 - :UNL1:O | 2.41198 | :UNL1:H5* - A:A34:N7 | 2.8529 |
| :UNL1:H - A:G4:O6 | 2.0637 | **192-5p-ERCC4-Ligand 11** | |
| **146a-5p-BRCA1-Ligand 1** | | A:A34:H62 - :UNL1:O | 2.74014 |
| A:C33:H41 - :UNL1:N | 2.4151 | :UNL1:H - A:U13:OP2 | 2.08923 |
| A:C33:H42 - :UNL1:O | 2.8326 | :UNL1:H - A:U13:O4 | 2.58961 |
| A:C34:H41 - :UNL1:O | 2.54852 | :UNL1:H - A:U13:O4 | 2.76051 |
| A:A35:H61 - :UNL1:O | 2.02418 | :UNL1:H - A:C14:OP2 | 2.40468 |
| **146a-5p-BRCA2-Ligand 11** | | **192-5p-ERCC4-Ligand 12** | |
| A:G29:HO2' - :UNL1:O | 2.63101 | A:C14:H42 - :UNL1:O | 2.84784 |
| A:A36:H61 - :UNL1:O | 2.55452 | :UNL1:H - A:G33:O6 | 2.40746 |
| A:A36:H62 - :UNL1:O | 2.56822 | :UNL1:H - A:A31:N7 | 2.23984 |

| | |
|---|---|
| A:A6:H62 - :UNL1:O | 2.46738 |
| :UNL1:H - A:U5:O4 | 2.53815 |
| :UNL1:H - A:A6:N7 | 2.05731 |
| **155-5p-MLH1-Ligand 11** | |
| A:A11:H61 - :UNL1:O | 2.07817 |
| A:A30:H61 - :UNL1:O | 1.8294 |
| A:A30:H62 - :UNL1:O | 2.59752 |
| :UNL1:H - A:U28:OP2 | 2.18114 |
| :UNL1:H - A:A30:N7 | 2.64405 |
| :UNL1:H - A:U31:O4 | 2.99175 |
| :UNL1:H - A:U28:O4 | 1.81047 |
| :UNL1:H - :UNL1:O | 2.56318 |
| **155-5p-MLH1-Ligand 12** | |
| A:A11:H61 - :UNL1:O | 2.13044 |
| A:A30:H61 - :UNL1:O | 2.41847 |
| A:A30:H62 - :UNL1:O | 2.52885 |
| :UNL1:H - A:A29:OP2 | 2.79143 |
| :UNL1:H - A:U28:O4 | 2.65692 |
| **155-5p-MLH1-Ligand 13** | |
| :UNL1:H - A:U31:O4 | 2.34373 |
| :UNL1:H - A:U31:O4 | 2.07935 |
| :UNL1:H - A:A29:OP2 | 2.1918 |
| :UNL1:H - A:C27:OP2 | 2.55713 |
| **155-5p-MSH6-Ligand 2** | |
| :UNL1:HN - A:G86:N7 | 2.36628 |
| :UNL1:HN - A:G85:O6 | 2.39277 |
| :UNL1:HN - A:G85:N7 | 2.3351 |
| :UNL1:H - A:U83:OP1 | 2.0833 |
| :UNL1:H - A:G86:N7 | 2.64568 |
| **155-5p-MSH6-Ligand 9** | |
| A:C29:H41 - :UNL1:O | 2.31302 |
| A:G55:H22 - :UNL1:F | 2.10075 |
| A:G55:H1 - :UNL1:F | 2.60674 |
| A:G56:H1 - :UNL1:O | 2.41582 |
| :UNL1:H - A:A31:OP2 | 2.5433 |
| :UNL1:H - A:U30:OP2 | 2.37947 |
| :UNL1:H - A:A31:OP2 | 2.71027 |
| :UNL1:H - A:G56:O6 | 2.276 |
| :UNL1:H - A:U32:O4 | 2.73291 |
| **192-5p-ERCC4-Ligand 13** | |
| A:A15:H61 - :UNL1:F | 2.2294 |
| :UNL1:H - A:U13:OP2 | 1.76569 |
| :UNL1:H - A:A12:OP1 | 2.96561 |
| :UNL1:H - :UNL1:O | 2.52992 |
| **203a-3p-ABL1-Ligand 4** | |
| A:A27:H61 - :UNL1:O | 2.20572 |
| A:C14:H41 - :UNL1:O | 2.74709 |
| **203a-3p-ABL1-Ligand 5** | |
| A:A27:H61 - :UNL1:O | 2.51662 |
| A:A27:H62 - :UNL1:O | 2.82307 |
| A:C14:H41 - :UNL1:O | 2.31441 |
| :UNL1:H - A:U16:O4 | 2.01305 |
| :UNL1:H - A:A15:N7 | 2.83259 |
| **203a-3p-ABL1-Ligand 6** | |
| A:A25:H61 - :UNL1:O | 2.16259 |
| A:A26:H61 - :UNL1:F | 2.66732 |
| A:A27:H61 - :UNL1:F | 2.7299 |
| A:C14:H42 - :UNL1:O | 2.80401 |
| :UNL1:H - A:A27:N7 | 2.97476 |
| :UNL1:H - A:G29:N7 | 2.62171 |
| :UNL1:H - A:U30:O4 | 2.24629 |
| **203a-3p-ATM-Ligand 1** | |
| A:U25:HO2' - :UNL1:O | 2.52847 |
| A:A27:H61 - :UNL1:O | 2.05028 |
| A:A28:H61 - :UNL1:O | 2.36792 |
| A:C21:H41 - :UNL1:O | 2.13329 |
| A:C21:H42 - :UNL1:N | 3.0013 |
| **203a-3p-ATM-Ligand 2** | |
| :UNL1:HN - A:U32:O4 | 2.37543 |
| :UNL1:HN - A:G31:OP2 | 2.60717 |
| :UNL1:H - A:G31:OP2 | 2.83116 |
| **203a-3p-ATM-Ligand 7** | |
| A:C16:H41 - :UNL1:O | 2.46751 |
| :UNL1:H - A:U14:OP2 | 2.27725 |
| :UNL1:H - A:G15:O6 | 2.2415 |
| :UNL1:H - A:G15:N7 | 2.74291 |

(*) Ligands having the best binding with complexes are ligands 2, 4, and 6. It can be noted that ligands 4, 5, and 6 are in fact isomers.
